# “I hear with my little ear something…": Enhanced performance of absolute pitch musicians in an interleaved melody recognition test

**DOI:** 10.1101/469429

**Authors:** Teresa Wenhart, Ye-Young Hwang, Eckart Altenmüller

## Abstract

Autistic people exhibit enhanced abilities to find and extract visual or auditory figures out of a meaningful whole (disembedding). Studies have shown heightened autistic traits in professional musicians with absolute pitch (AP). This study investigates whether such musicians show an advantage in an interleaved melody recognition task (IMRT).

A total of N=59 professional musicians (AP=27) participated in the study. In each trial a probe melody was followed by an interleaved sequence. Subjects had to indicate as to whether the probe melody was present in the interleaved sequence. Sensitivity index d’ and response bias c were calculated according to signal detection theory. Additionally, a pitch adjustment test measuring fine-graded differences in absolute pitch proficiency, the Autism-Spectrum-Quotient and a visual embedded figures test were conducted.

AP performance was enhanced overall compared to RP. Absolute pitch proficiency, visual disembedding ability and musicality predicted approximately 39.2% of variance in the interleaved melody recognition test. No correlations were found between IMRT and autistic traits.

The stable pitch-label associations of AP might serve as additional sensory cues during pre-attentive processing in recognizing interleaved melodies. Results are in line with a detailed-oriented cognitive style and enhanced perceptional functioning of AP musicians similar to that observed in autism.

## Background

One of the most complex functions of the auditory system lays in the ability to disentangle and extract distinct streams (“auditory streaming”) of information out of the mixture of sounds that reach the ear (e.g. two interleaved and overlapped melodies played by different instruments in an orchestra or isolating a voice among noise in a crowded room). This is a key component of “auditory scene analysis”^1^ with two different mechanisms: pre-attentive so-called “primitive” (bottom up) processes and, often but not always attentive, “schema-based” (top down) mechanisms^1^. Primitive processes sequentially or simultaneously separate the incoming sound mixture based on sensory information, i.e. according to regularities of harmonicity, or loudness, or in relation to spatial orientation ^1^, or according to similarity or smooth changes in pitch^2–4^, timbre^3^, rhythm^5^, or meter^6^ and seem to be innate^7^. These factors roughly correspond to the famous Gestalt principles^8^ of e.g. similarity, proximity and many others that have reached wide acceptance among visual perception researchers. On top of that, music or speech schemas learned via experience are used to attentively compare, extract and structure the auditory environment based on previous acquired knowledge^4,6^. Auditory streaming has been investigated using an interleaved melody recognition task, first developed by W.J. Dowling (1973)^4^ and later adapted and extended by Bey and McAdams (2002, 2003)^4,6^ or unknown melody^2,3^ is presented first, followed by an interleaved sequence, in which either the target or a modified melody is interleaved with distractor tones in the same pitch range.

It is not surprising that musicians exhibit an enhanced ability to extract musical streams out of complex interleaved melodies^3^, as they belong to a population with far greater than average hearing abilities and knowledge of music theory^9,10^. However, it remains to be investigated, if absolute pitch (AP) possessors, who have the unique (<1%)^11^ ability to name or produce a musical tone without a reference^12,13^, differ in auditory streaming from relative pitch possessors. Absolute pitch also occurs far more often in professional musicians (7-15 %)^14–16^ than in the general population, but it is still unclear why few people develop absolute pitch while most humans are relative pitch possessors. Among possible predictive factors for AP are an early onset of musical training especially before the age of 7 ^16,17^ and related to this a critical period^18–22^, genetic contributions^18^, the type of musical education method^16^ and ethnicity^14,15,23^. Most likely, absolute pitch is the result of an interaction between genes and environmental influences, which makes the ability a fascinating research topic for human cognitive neuroscience^22^.

The ability of absolute pitch possessors to not only judge pitch heights but also perceive an additional quality for different tone categories, so-called pitch “chroma”, or by contrast the ability of relative pitch (RP) possessors to neglect single note information but focus on melodically more relevant information of musical intervals, might lead to performance differences in experiments on auditory streaming.

Furthermore, the indirect measurement of auditory stream segregation with the use of interleaved melody recognition tests is very similar to the concept of disembedding in autism research and has indeed been described as an “auditory hidden figures test” by Dowling, Lung & Herrbold (1987)^6^. It has been shown, that people with autism-spectrum-conditions (ASC) show enhanced abilities to extract small visual or auditory figures out of a meaningful whole respectively a fully interleaved auditory sequence^23–27^. However, autistic people seem to fail to use pitch separation cues in auditory streaming experiments^23,25^.

Autism-spectrum-conditions, which are characterized by social and communication difficulties, sensory abnormalities, narrow interests and problems with unexpected change^28^, sometimes co-occur with special abilities^29–32^, one of the more frequent of which is absolute pitch^31,33–35^. Finally, recent research has shown increased autistic traits in absolute pitch possessors^36,37^. It has been suggested, that this joint occurrence might be explained by similar brain network structure and function^31,38–40^ and a detail-oriented cognitive style^19,31^ among people of those populations. However, it is still a matter of debate (a) whether absolute pitch possessors also exhibit an enhanced ability for disembedding in vision and audition (auditory streaming) similar to autism, (b) whether this is correlated to autistic traits in AP possessors and (c) whether having absolute pitch might explain different strategies and performance during auditory streaming experiments. The present study aims to shed new light onto the discussion by investigating professional musicians with and without absolute pitch as they take part in an auditory streaming experiment created after Bey & McAdams^2,3^ and an visual embedded figures test. We also determine their scores for autistic traits. We hypothesize that the ability of auditory stream segregation can be predicted by absolute pitch proficiency, autistic traits and musicality and reflects an auditory equivalent of disembedding in vision.

## Methods

This study was formed part of a larger project consisting of various experiments investigating cognitive performance in audition and vision of absolute vs. relative pitch possessors. Therefore parts of the methods (sample description, general procedure, description of absolute pitch assessment, covariate measurements) are by their nature similar to previous publications of our lab (Wenhart, Bethlehem, Baron-Cohen & Altenmüller, *under review*; Wenhart, & Altenmüller, *under review*).

### Participants

Thirty-one AP musicians (16 female) and 33 RP musicians (15 female) - primarily students or professional musicians at the University for Music, Drama and Media, Hanover - were recruited via an online survey using UNIPARK software (https://www.unipark.com/). Four AP and two RP were amateur musicians. Non-native German speakers (4 AP) had the choice between a German and an English version of the experiments. All participants denied any history of severe psychiatric or neurological conditions and one AP reported taking Mirtazapine. The primary instruments played by the AP were piano (15), string instruments (9), woodwind instruments (3), voice (2), and brass instruments (2); for RP they were piano (13), string instruments (4), woodwind instruments (6), voice (3), brass instruments (3), bass (1), guitar (1), accordion (1), and drums (1). All AP but one were consistently right handed according to the Edinburg Handedness Iventory^41^; three RP were left-handed, two RP were ambidextrous. A total of five participants (4 AP, 1 RP; final sample: AP=27, RP=32) were excluded during analysis due to missing data (1 case), extreme values for reaction times (2 cases) or because they did not follow the instructions (2 cases). The study was approved according to Helsinki Declaration by the ethic committee of the Hanover Medical School (Approval no. 7372, committee’s reference number: DE 9515). The methods were carried out in accordance with their guidelines and regulations. All participants gave written consent.

### General procedure

The overall project consisted of three parts: one online survey and two appointments in the lab of the Institute for Music Physiology and Musicians’ Medicine, Hanover. The online survey was used for demographic questions, Autism Spectrum Quotient (AQ^42^, German version by C.M. Freiburg, available online: https://www.autismresearchcentre.com/arc_tests), Musical Sophistication Index (GOLD-MSI^43^ (questionnaire)), an in-house questionnaire assessing musical education as well as total hours of musical training within the life span and a preliminary assessment of absolute pitch ability in order to allocate subjects to groups (AP vs. RP). The absolute pitch assessment was performed using an in-house pitch identification screening (PIS) consisting of 36 categorical, equal-tempered sine waves in the range of three octaves between C4 (261.63 Hz) and B6 (1975.5 Hz). Subjects exceeding 33% correct trials (>12/36 tones named correctly) were assigned to the AP group. Tests on general intelligence (Raven’s Standard Progressive Matrices (SPM))^44^, information processing speed (“Zahlenverbindungstest” (ZVT))^45^, a musical ability test (Advanced Measures of Music Audiation (AMMA))^46^, a visual Embedded Figures Test (Group Embedded Figures Test (GEFT))^47^ and all experiments were conducted in the lab (see Table 1 for general group differences regarding covariates).

**Table 1.** Participants’ characteristics

|  | AP (n=27) |  |  | RP (n=32) |  |  | t-test<br>(two-tailed) |
| --- | --- | --- | --- | --- | --- | --- | --- |
|  | Mean | SD | Range | Mean | SD | Range |  |
| Age | 25.7 | 9.7 | 17-58 | 24.0 | 7.13 | 17-57 | $t(46.9) = -0.756$;<br>$p = 0.453$ |
| SPM-IQ | 114.67 | 12.59 | 74.5-132.25 | 114.2 | 13.3 | 86.5-134.5 | $t(56.2) = -0.138$;<br>$p = 0.891$ |
| ZVT-IQ | 123 | 12.47 | 101.5-145 | 120.5 | 13.89 | 97-143.5 | $t(56.7) = -0.728$;<br>$p = 0.470$ |
| Hours main instrument | 11551.27 | 9540.95 | 1642.5-39785 | 14107.82 | 17263.78 | 1606-77617.25 | $t(49.7) = 0.437$;<br>$p = 0.664$ |
| AMMA | 65.59 | 6.04 | 53-78 | 63.28 | 7.14 | 46-76 | $t(57) = -1.348$;<br>$p = 0.183$ |
| MSI | 209.04 | 14.79 | 182-234 | 211.12 | 15.24 | 185-246 | $t(55.8) = 0.533$;<br>$p = 0.596$ |
| **PIS** | 28.92 | 6.25 | 15-36 | 5.44 | 4.33 | 0-21* | **$t(43.1) = -16.25$;**<br>**$p < 2.2e-16$** |
| **AQ** | 20.11 | 6.23 | 10-36 | 16.88 | 5.52 | 6-27 | **$t(52.5) = -2.093$;**<br>**$p = 0.041$** |
| **MAD** | 42.16 | 39.02 | 9.8 -200.57 | 297.77 | 87.33 | 91.04 -467.52 | **$t(44.4) = 14.890$;**<br>**$p < 2.2e-16$** |
| **SDfoM** | 51.57 | 47.27 | 7.41-235.69 | 330.04 | 124.72 | 134.37 -811.73 | **$t(41.0) = 11.675$;**<br>**$p = 1.272e-14$** |
| **Starting age** | 5.56 | 2.67 | 2-17 | 7.19 | 2.22 | 3-12 | **$t(55.8) = 2.527$;**<br>**$p = 0.015$** |

Age, nonverbal IQ (SPM), information processing capacity (ZVT), musical training (total hours during life span on main instrument), musicality (AMMA; MSI) and online pitch identification screening (PIS) for each group; \* two RP reported not having absolute pitch but reached a screening score of 13 respectively 21. Because of this and their weak performance in the pitch adjustment test, the subjects were assigned to the RP group; Significant group differences highlighted in bold. $p < .10$ , $p < .05$ \*, $p < .01$ \*\*, $p < .001$ \*\*\* (uncorrected)

### Experiments and material

#### Pitch adjustment test (PAT)

A pitch adjustment test (PAT) developed after Dohn et al. ^48^ was used to quantify fine-graded differences in absolute pitch proficiency. The test consisted of 108 target notes, presented as letters on a PC screen in semi-random order in 3 Blocks of 36 notes each (3*12 different notes per block) with individual breaks between the blocks. Participants’ task was to adjust the frequency of a sine wave with random start frequency (220 - 880 Hz, 1Hz steps) and to try to hit the target note (letter presented centrally on PC screen, e.g. “F# / Gb”) within at most 15 seconds. Participants were allowed to choose their octave of preference. The tones were presented through sound isolating Shure 2-Way-In-ear Stereo Earphones (Shure SE425-CL, Shure Distribution GmbH, Eppingen, Germany). Subjects were explicitly asked to try to adjust each tone as precisely as possible without the use of any kind of reference and to confirm their answer with a button press on a Cedrus Response Pad (Response Pad RB-844, Cedrus Corporation, San Pedro, CA 90734, USA). If no button was hit, the final frequency after 15 seconds was taken and the experiment proceeded with the next trial. In both cases, the Inter Trial Interval (ITI) was set to 3000ms. Online pitch modulation was performed by turning a USB-Controller (Griffin PowerMate NA16029, Griffin Technology, 6001 Oak Canyon, Irvine, CA, USA) and implemented in Python according to Dohn et al. ^48^. Participants could choose between rough (10 cent, by scrolling the wheel) and fine tuning (1 cent, by pressing down and scrolling the wheel). The final or chosen frequencies of each participant were compared to the nearest target tone (< 6 semitones/600cent). For each participant, mean absolute deviation (MAD^48^, equation (1)) from target tone

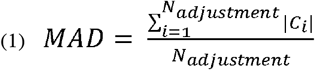

was calculated as the mean of the average absolute deviations c_i_ of the final frequencies to the target tone (referenced to a 440 Hz equal tempered tuning).

MAD reflects the pitch adjustment accuracy of the participants. The consistency of pitch adjustments (SDfoM, standard deviation from own mean), possibly reflecting the tuning of the pitch template^48^, was then estimated by taking the standard deviation of the absolute deviations (equation (2)).

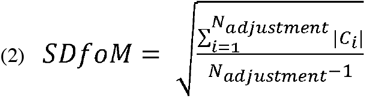

For regression analyses (see below), we performed a z-standardization of the MAD (Z_MAD, equation (3)) and SDfoM (Z_SDfoM, equation (4)) values relative to the mean and standard deviation of the non-AP-group, as originally proposed by Dohn et al. (2014)^48^.

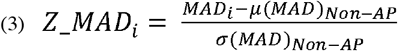

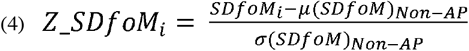

#### Interleaved Melody recognition test (IMRT)

The experiment consisted of 16 training trials followed by 128 test trials presented in two blocks of 64 trials each with a break in between. Each trial was made up of a 6-tone probe melody (2400 ms, 400ms per tone; ~74.5bpm) followed by an interleaved sequence (see figure 1), in which the probe melody was embedded within a distractor melody (1^st^ and then every 2^nd^ tone belonging to the probe melody; 12 tones, 2400ms, 200ms per tone). In half of the trials, a modified version of the probe melody was embedded within the interleaved sequence. The general task for the participants was to press the right button on the Cedrus Response Pad (Response Pad RB-844, Cedrus Corporation, San Pedro, CA 90734, USA), if the probe melody was embedded inside the interleaved sequence, and the left button if not (required responses were counterbalanced; button colors (yellow and blue) were counterbalanced across subjects). Each trial was preceded by the German or English word for “Attention!” lasting for 1000ms, and probe melody and interleaved sequence were separated by an ISI (Inter-Stimulus-Interval) of 1000ms. The ITI (Inter-Trial-Interval) was set to 2200ms.

**Figure 1.**
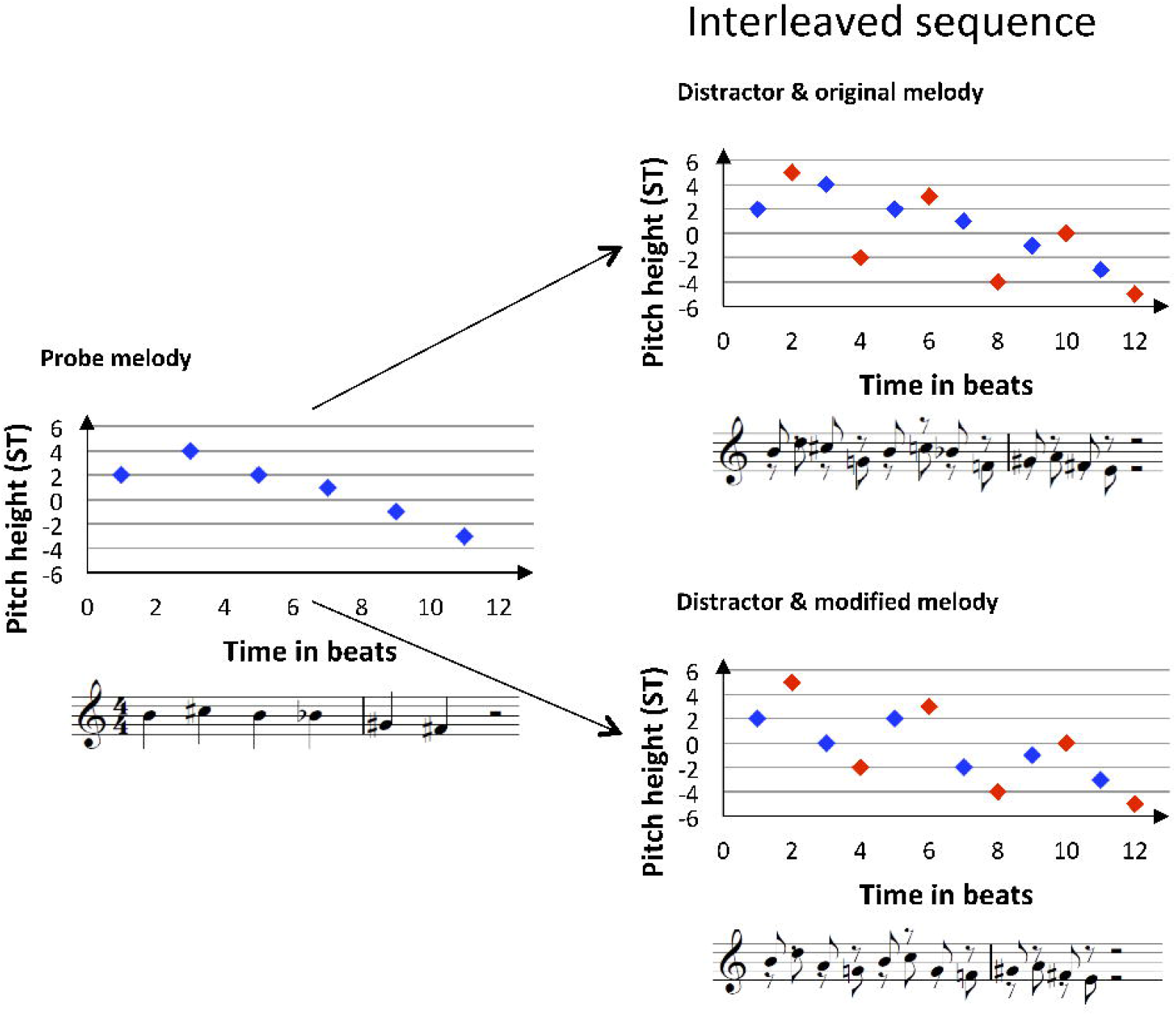
Graphical and musical notation of experimental material within the IMRT (interleaved melody recognition test). Each trial begins with a probe melody (A, blue) followed by an interleaved sequence with either the same probe melody (B, blue) or a modified melody (C, blue, here 2^nd^ and 4^th^ note modified compared to probe melody) interleaved with a distractor melody (red). Graphical notation of musical melodies in this example shows the pitch height of each note in ST relative to the A_4_ (440 Hz). The distractor notes always encompassed the neighboring melody notes in terms of the pitch height.

Probe melodies, modified probe melodies and distractor melodies were constructed according to Bey & McAdams (2002, 2003)^2,3^ as follows:

The mean frequency of each probe melody was within a range from −3 to +2 semitones (ST) around the equal-tempered A_4_ (440 Hz), i.e. from F^#^_4_ to B_4_. Modified melodies were then created by altering the 2^nd^ and 4^th^ or the 3^rd^ and 5^th^ note of the probe melody within a range of +-4 ST, which always also altered the melodic contour (see Appendix A of Bey & McAdams (2003)^3^). All melodies ranged from 5 to 11 ST and intervals within the melodies were between 1 and 8 ST. The 36 probe melodies and 36 modified melodies comprised 46 diatonic and 26 non-diatonic melodies. For each of the 36 melody pairs (probe and modified version) one distractor melody was composed (see Appendix B of Bey & McAdams (2003)^3^). All notes of distractor melodies were maximally 1-2 ST (alternating) above respectively below the two neighboring tones of the target melody to ensure, that the task could not be solved by the global contour of the sequence. In half of the melodies the first distractor note started below the target melody and the reverse was true for the other half. Finally, 8 interleaved sequences were built for each of the 36 melody pairs: distractor and either probe or modified melody were interleaved and – relative to the note closest to the mean frequency - separated by 0 (fully interleaved), 6 (separated by augmented fourth), 12 (one octave) or 24 ST (two octaves, see figure 2).

**Figure 2.**
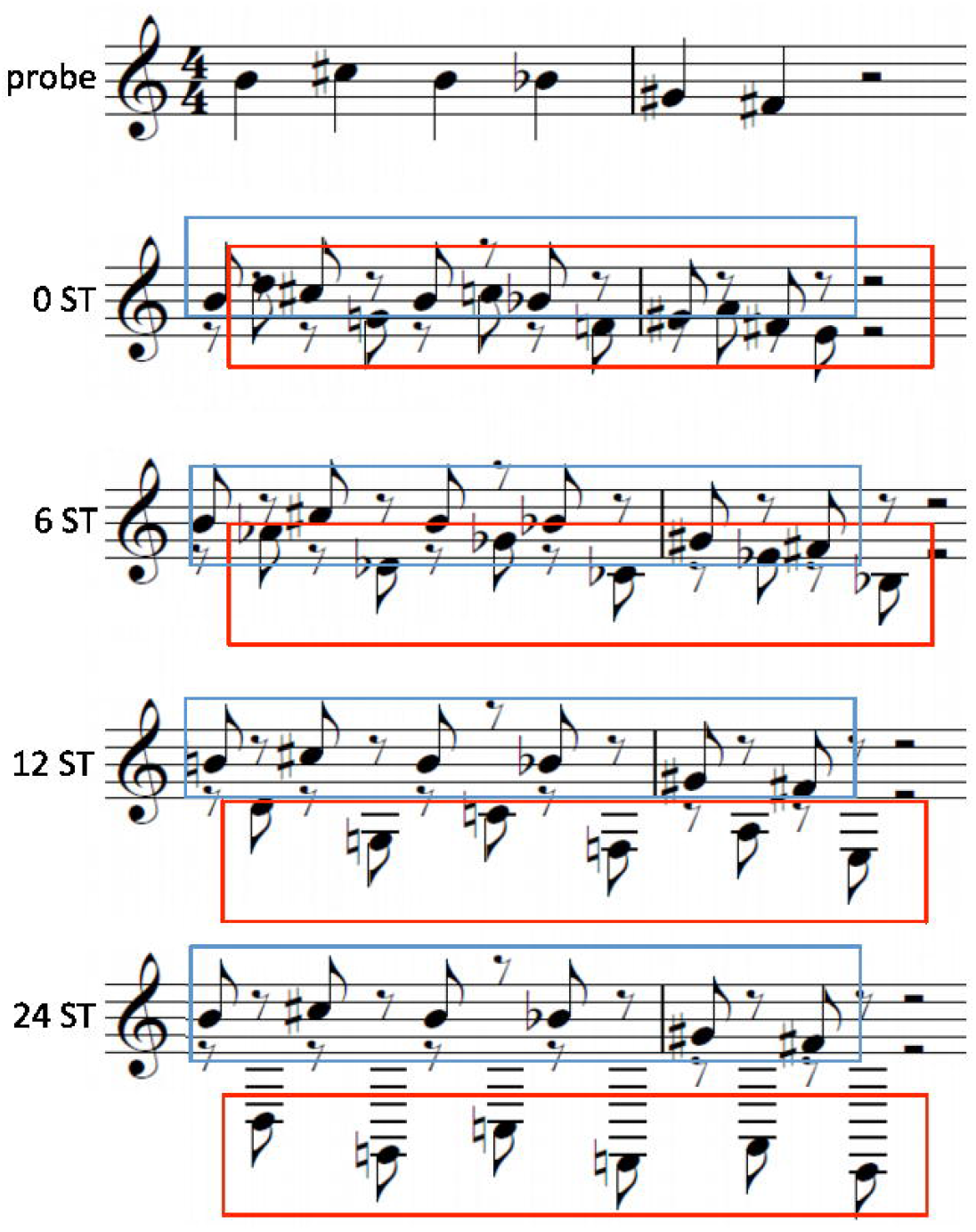
Interleaved sequences of IMRT. Probe melody respectively modified probe melody (here probe melody, blue) and distractor melody (red) were interleaved and four separation conditions created. The two melodies were either fully interleaved and their mean frequency separated by 0 semitones (0 ST) or the distractor melody was shifted by 6, 12 or 24 ST downwards. Therefore melodies were sequentially separated further apart, increasing the possibility of hearing two musical streams.

Two experimental versions (A, B) were created, with half of the participants receiving the trials related to probe melody 1-18 and the other half receiving the trails related to probe melodies 19-36. The 16 trials of the first two (Version A), respectively 19^th^ and 20^th^ target melodies (Version B) served as training trails. Order of trials was pseudo-randomized with the same probe melody never occurring twice in a row. The experiment was programmed and controlled via the Python toolbox PsychoPy^49,50^. Due to a technical error in the first version, one trial occasionally appeared twice during the experiment while another one was missed. The error occurred randomly with respect to probe vs. modified melody and semitone separation. To control for possible effects on the statistical tests, an equal number of participants in each group performed this imperfect version (N=12; final percentage after exclusions: RP: 12/31, AP: 9/27), and all other participants afterwards performed the correct version. Calculation of Hit rates and False alarm rates was adapted accordingly, as described below. The total experiment without instructions lasted approximately 21.6 minutes.

#### Group Embedded Figures Task (GEFT)

The group embedded figures test (GEFT)^47^, a paradigm often used to investigate cognitive style in autism^24,25,51^, was chosen to additionally assess subjects ability to find smaller shapes in drawings of a meaningful whole. The test is performed with pen and paper and consists of 18 trials in two blocks of 9. The participant has to find a geometric form in a large and more complex whole and mark it with a pencil. As we observed ceiling effects as to the total amount of correct trials in the GEFT during pilot experiments, we additionally measured the time subjects needed to find each form so that we could better discriminate between subjects’ abilities at this task. Performance on the GEFT was therefore expressed as average time (over correct trials) needed to solve an item.

### Statistical Analysis

All statistical analyses were performed using the open-source statistical software package R (Version 3.5, https://www.r-project.org/). As melodies unfold over time, no reaction times were analyzed with respect to the IMRT. Instead, sensitivity index *d`* and response bias (decision criterion) *c* were calculated according to signal detection theory^52,53^. Signal detection theory is a suitable method for analyzing decision processes in recognition or discrimination tasks like in the IMRT, in which only two responses are possible (here: whether or not the probe melody is embedded). Therefore, responses fall into one of four categories: hit (correct identification of probe melody in the interleaved sequence), miss (miss of the probe melody in the interleaved sequence), false alarm (false “yes” response to modified melody in the interleaved sequence) or correct rejection (correct “no” response to modified melody in the interleaved sequence). According to Macmillan^54^ the difference of the *z*-scores of proportions of hits (H in %) and false alarms (F in %) gives the sensitivity index *d`*, which is independent of the decision criterion *c*.

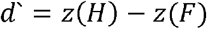

The latter is a measure of the tendency to say “yes” or “no”, i.e. the response bias, and is calculated as:

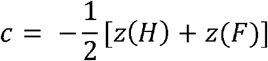

Therefore, positive values of *c* indicate a tendency towards “no” responses, while negative values indicate a tendency towards “yes” responses. As *z*-values for perfect proportions yielding H=1 and F=0 cannot be calculated (infinite values), Macmillan ^54^ recommends to reduce the proportion of N_correct_/N_total_ by subtracting 0.5 point from the number of correct trials within the affected condition: H = (N_correct_-0.5)/N_total,_ respectively to increase the proportion of false alarms by adding 0.5 points: F = (N_false_+0.5)/N_total_. The IMRT comprised (in the errorless version) 16 trials per condition (8 conditions: 4 separation conditions x 2 melody conditions (probe/modified)); with this number of trials, some perfect scores were likely. As 21 subjects had unequal trial numbers per condition (see figure A.1, Supplementary Material A) hit rates and false alarm rates were always calculated according to the true number of trials of each subject per condition. However, since different numbers of trials affect calculated proportions and *z*-values, we performed additional calculations (e.g. using the number of trials of the subject with the most trials per condition as N_total_) of proportions for hit rates and false alarm rates. These alternative analyses did not affect the direction or effect sizes of the reported statistical tests.

## Results

### IMRT group differences

Statistical tests were based on the signal detection measures mentioned above, with perceptual sensitivity (d)’ and “response bias” (c) inspected separately. The two versions of IMRT (A, B), each containing half of the melodies, unintentionally differed in terms of d’ scores. On average, version A yielded lower scores than version B (see table B1 Supplementary Material B). Therefore, a dichotomous variable containing information of version performed per subject was included in the analyses as a covariate. In contrast, different numbers of trials within frequency separation conditions (see Methods section) did not lead to significantly different results either in overall performance or for separate conditions (see table A.1, Supplementary Material A for further details), so number of trials was not included as a covariate.

First and importantly, the validity of IMRT (hence d’ values) for measuring disembedding in audition was confirmed by Pearson correlation analysis (r = −0.407, *p* =0.0014) with reaction times (RT in s) on the visual embedded figures test (GEFT)^47^. In total, roughly 15 % of variance in the overall performance on IMRT was explained by the average time needed to solve the items of GEFT (linear model: intercept: 4.096 ***, GEFT: −0.042 **; F (1, 57) = 11.28, *p* = 0.0014; R^2^ = 0.165, R^2^_adjusted_ = 0.151; see figure 3A).

**Figure 3.**
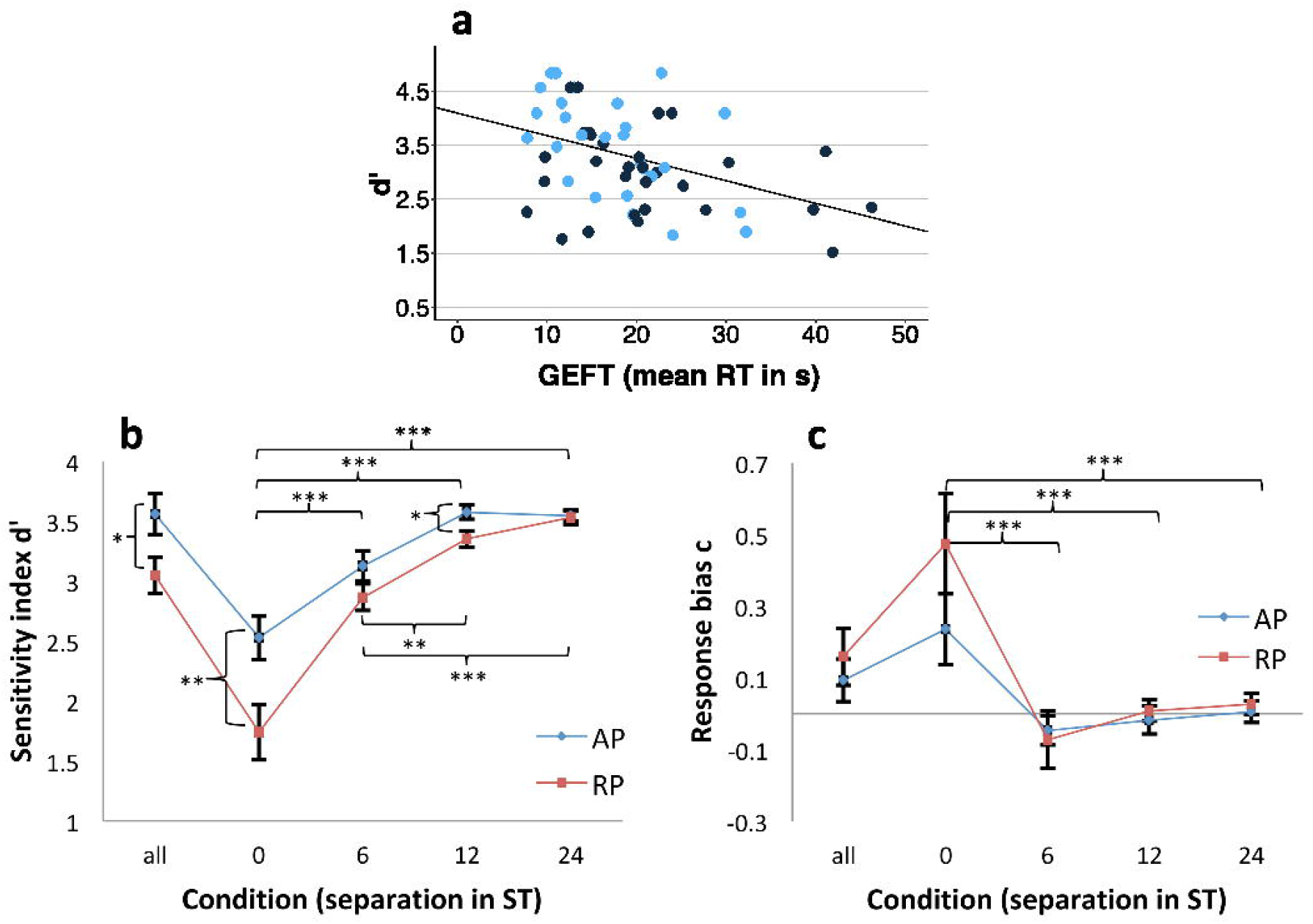
Relation of IMRT and visual embedded figures test (a) and signal detection measures by group (b, c). (a) Overall performance on IMRT (d’) by mean time (in s) needed to solve an item on the visual embedded figures task (GEFT, ^47^). Variables revealed a correlation of r = −0.407 (Pearson correlation; *p* <.001). Blue: IMRT version A, black: IMRT version B. (b, c) Results on sensitivity index d` (b) **and** response bias c (^54^ by group (AP, absolute pitch; RP, relative pitch) regarding overall performance (“all”) and performance within separation conditions (0, 6, 12, 24 semitone (ST) separations) on IMRT. Higher values of d’ indicate better performance. Positive values of c indicate a tendency towards “no” responses, negative values a tendency towards “yes” responses. Bars represent standard errors. * p<.05. ** p<.01. *** p<.001.

Hence, IMRT was deemed to be an adequate test for investigating auditory disembedding. Therefore, a 2×4 repeated measurements ANOVA with separation (0, 6, 12, and 24) as within-subject factor and group (AP vs. RP) as between subject factor was run. Analyses revealed main effects of group (F (1, 59) = 5.901, *p* = 0.0183, η^2^_partial_ = 0.138) and separation (F (4, 228) = 76.474, *p* = 2e-16, η^2^_partial_ = 0.573) as well as a significant interaction (F (4, 228) = 4.898, *p* = 0.000831, η^2^partial = 0.079; see figure 3B).

Post hoc t-tests revealed significant differences between the 0-ST-condition and all other separation conditions (6-ST: *p* = 2e-16, 12-ST: *p* = 2e-16, 24-ST: *p* = 6.3e-9) as well as between the 6-ST-condition on the one hand and both the 12-ST-(*p* = 0.00452) and 24-ST-conditions (*p* = 0.00058), see figure 3B) on the other hand. Furthermore, absolute pitch possessors performed significantly better on the 0-ST and 12-ST conditions and when considering overall performance (see table 2). We would also like to draw attention to a likely ceiling effect in condition 24-ST (see figure 3B and figure 4E), the results of which should be interpreted with caution.

**Figure 4.**
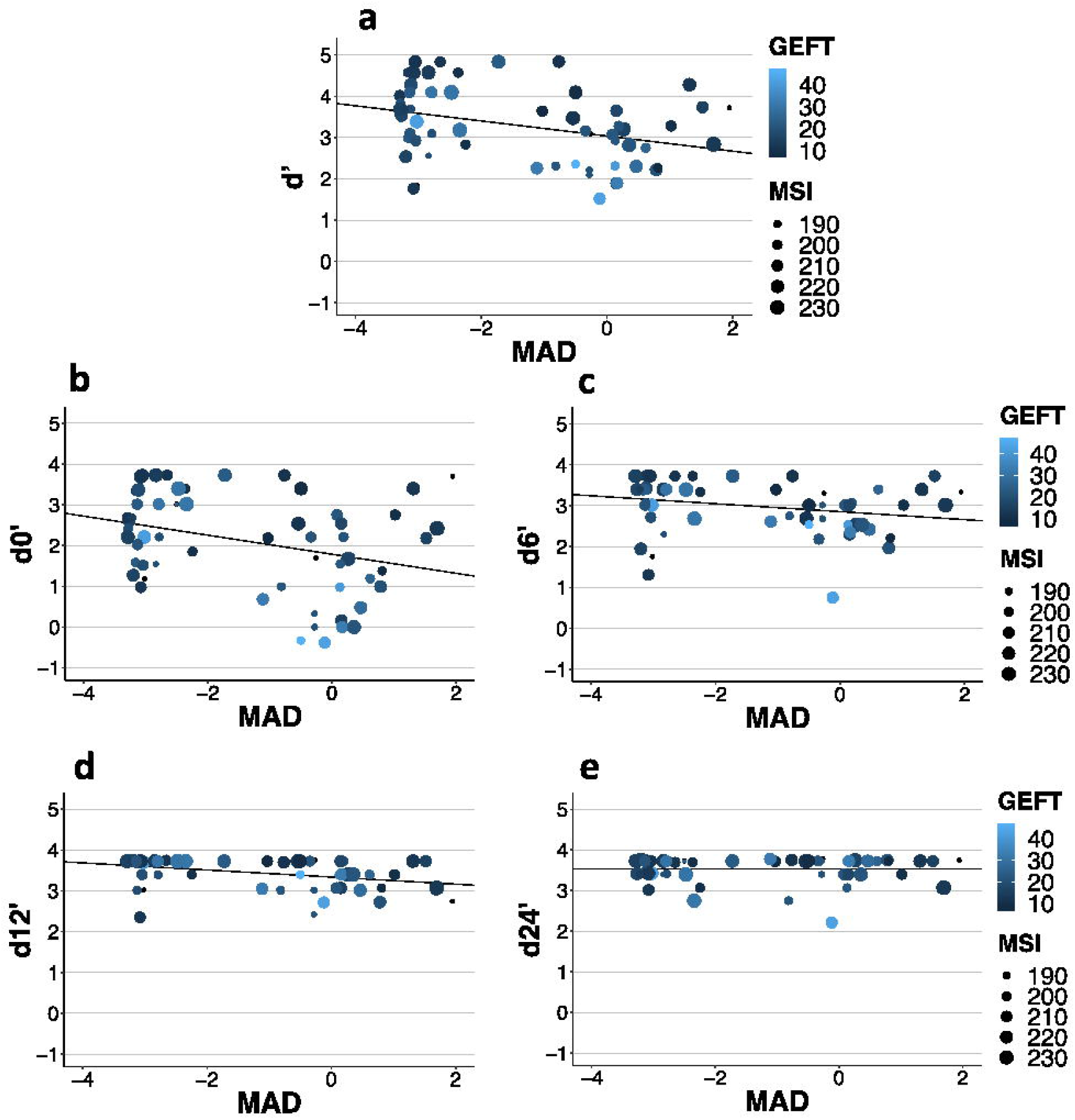
Influence of absolute pitch ability (Z_MAD), disembedding in vision (GEFT, time in s) and musicality (MSI) on perceptual sensitivity d’. Panel A corresponds to overall performance on IMRT, while panels B-E show the prediction of performance in different separation conditions (0, 6, 12, 24 semitone separation of probe and distractor melody). Color and shape scales correspond to disembedding in vision (GEFT, time in s) respectively MSI (score on questionnaire, higher values indicate greater musical sophistication). The regression line always takes the intercept and beta-weight of the simple linear regression of absolute pitch ability (Z_MAD, standardized to mean of RP group) on sensitivity index (d’-d24’, higher values indicate better performance). Ceiling effects in the 12-ST- and especially 24-ST-condition (panel E) are clearly visible (see table 4 for statistical values).

**Table 2.** IMRT group differences (N=59)

|  | AP |  | RP |  | t-test |  |  |
| --- | --- | --- | --- | --- | --- | --- | --- |
|  | Mean | SD | Mean | SD | t (df) | p-value | Cohen's d |
| All | 3.559 | 0.868 | 3.045 | 0.872 | -2.262(55.4) | < .028 * | -0.591 |
| 0 ST | 2.527 | 0.923 | 1.739 | 1.279 | -2,739 (55.8) | < .008 ** | -0.697 |
| 6 ST | 3.127 | 0.650 | 2.865 | 0.620 | -1.565 (54.4) | 0.123 | -0.411 |
| 12 ST | 3.575 | 0.340 | 3.350 | 0.380 | -2.411 (56.8) | < .019 * | -0.624 |
| 24 ST | 3.548 | 0.250 | 3.528 | 0.360 | -0,250 (55.3) | 0.804 | -0.063 |

Means, standard deviation (SD) and statistical comparison (post hoc t-test, two-tailed) of IMRT performance (perceptual sensitivity index d') by group (absolute pitch, AP, vs. relative pitch, RP) and per separation condition (in semitones, ST). \* p <.05, \*\* p<.01

With respect to response bias (c) no significant group differences were found for overall performance nor for separation conditions (see figure 3C and table B2, Supplementary Material B). However, a 2x 4 repeated measures ANOVA with separation (0, 6, 12, 24) as within-subject factor and group (AP vs. RP) as between subject factor revealed a significant main effect of separation (F (4, 228) = 15.594, *p* = 2.71e-11, η^2^_partial_ = 0.215). No main effect of group (F (1, 57) = 0.859, *p* = 0.358, η^2^_partial_ = 0.013) and no interaction of group and separation (F (4, 228) = 1.452, *p* = 0.218, η^2^_partial_ = 0.025) were found. Post hoc tests revealed a significant higher tendency to respond “yes” on 0-ST-trials compared to all other conditions (6-ST: *p* = 9.4e-6; 12-ST: *p* =3.2e-5; 24-ST: *p* = 2.0e-7). This again confirms the impression that a greater separation between probe and distractor melody decreased the difficulty of the task and hence led to better perceptual sensitivity (d’) and nearly no response bias (c, see figures 3B-C and 4).

### Linear models to predict perceptual sensitivity d’

To investigate which musical and cognitive variables influence performance on IMRT, we performed multiple linear regressions separately for overall performance and separation conditions (0, 6, 12, 24 ST). In a first step, bivariate correlations between IMRT and variables of interest were calculated to get an overview of possibly predictive variables (see table 3).

**Table 3.**
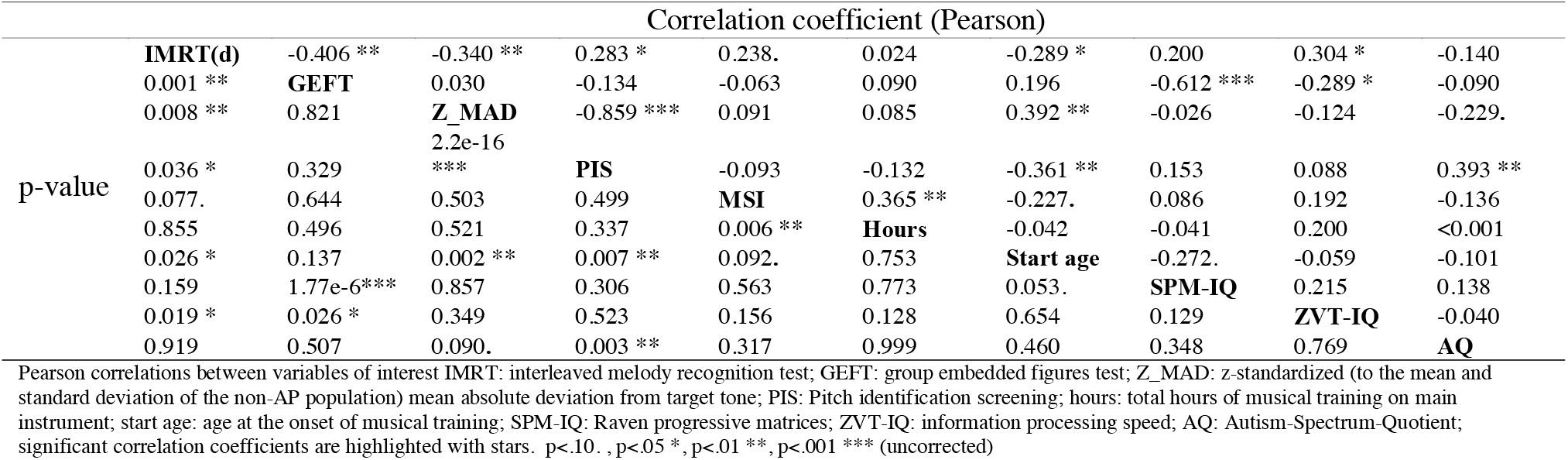
Bivariate correlations between variables of interest

| Correlation coefficient (Pearson) |  |  |  |  |  |  |  |  |  |  |
| --- | --- | --- | --- | --- | --- | --- | --- | --- | --- | --- |
| p-value | IMRT(d) | -0.406 ** | -0.340 ** | 0.283 * | 0.238. | 0.024 | -0.289 * | 0.200 | 0.304 * | -0.140 |
|  | 0.001 ** | GEFT | 0.030 | -0.134 | -0.063 | 0.090 | 0.196 | -0.612 *** | -0.289 * | -0.090 |
|  | 0.008 ** | 0.821 | Z_MAD | -0.859 *** | 0.091 | 0.085 | 0.392 ** | -0.026 | -0.124 | -0.229. |
|  |  |  | 2.2e-16 |  |  |  |  |  |  |  |
|  | 0.036 * | 0.329 | *** | PIS | -0.093 | -0.132 | -0.361 ** | 0.153 | 0.088 | 0.393 ** |
|  | 0.077. | 0.644 | 0.503 | 0.499 | MSI | 0.365 ** | -0.227. | 0.086 | 0.192 | -0.136 |
|  | 0.855 | 0.496 | 0.521 | 0.337 | 0.006 ** | Hours | -0.042 | -0.041 | 0.200 | <0.001 |
|  | 0.026 * | 0.137 | 0.002 ** | 0.007 ** | 0.092. | 0.753 | Start age | -0.272. | -0.059 | -0.101 |
|  | 0.159 | 1.77e-6*** | 0.857 | 0.306 | 0.563 | 0.773 | 0.053. | SPM-IQ | 0.215 | 0.138 |
|  | 0.019 * | 0.026 * | 0.349 | 0.523 | 0.156 | 0.128 | 0.654 | 0.129 | ZVT-IQ | -0.040 |
|  | 0.919 | 0.507 | 0.090. | 0.003 ** | 0.317 | 0.999 | 0.460 | 0.348 | 0.769 | AQ |

Pearson correlations between variables of interest IMRT: interleaved melody recognition test; GEFT: group embedded figures test; Z\_MAD: z-standardized (to the mean and standard deviation of the non-AP population) mean absolute deviation from target tone; PIS: Pitch identification screening; hours: total hours of musical training on main instrument; start age: age at the onset of musical training; SPM-IQ: Raven progressive matrices; ZVT-IQ: information processing speed; AQ: Autism-Spectrum-Quotient; significant correlation coefficients are highlighted with stars. p<.10. , p<.05 \*, p<.01 \*\*, p<.001 \*\*\* (uncorrected)

Apart from the already known correlation with GEFT (r = −0.406, *p* =0.0014), IMRT (overall performance) was related to absolute pitch proficiency in both producing a tone (Z_MAD, standardized mean absolute deviation from target tone; r = −0.340, *p* = 0.0085) and naming a tone (PIS, Pitch identification screening; r = 0.283, *p* = 0.0364), age of onset of musical training (r = −0.289, *p* = 0.0264), information processing speed (r = 0.304, *p* = 0.0199) and marginally to musical sophistication index (MSI; r = 0.238, *p* = 0.077).

For multicollinearity reasons age of onset and PIS were not included into regression models, as they showed high correlations with absolute pitch performance (Z-MAD: r = 0.392, *p* = 0.0021; PIS: r = −0.361, *p* = 0.00679) respectively pitch adjustment (Z_MAD: r = −0.859, *p* = 2.2e-16). Therefore a total of 5 variables (Z_MAD, MSI, GEFT, ZVT and version) were included into the 5 resulting regression models. Afterwards, variables with non-significant beta-weights were removed from the model leading to 5 reduced models (see table 4). In general IMRT performance was highly predicted by absolute pitch ability (Z-MAD) and GEFT. Performance in the 0-ST condition was additionally predicted by musical sophistication index (MSI) and the version of IMRT (A vs. B). Models of the 24-ST-condition did not reach significance and must be interpreted with caution due to ceiling effects (see figure 3B-C and figure 4E). From Figure 4 it can easily be seen that the four IMRT separation conditions (0, 6, 12, 24) decrease in variance (y-axis), indicative of decreasing task difficulty. Because at six semitones and greater separations the MSI is no longer predictive of IMRT (d’), this effect likely is due to our highly over-trained sample of professional musicians.

**Table 4.** Comparison of models predicting IMRT

| d' | Predictors (β) |  |  |  |  |  | comparison of models |  |  |  |
| --- | --- | --- | --- | --- | --- | --- | --- | --- | --- | --- |
|  | intercept | Z_MAD | MSI | GEFT | ZVT | version | F(df) | p-value | R<sup>2</sup> | R<sup>2</sup><sub>adjusted</sub> |
| full | -0.286 | -0.214 *** | 0.013. | -0.027 * | 0.007 | 0.447 * | 8.004 (5.5) | 1.33e-5 *** | 0.445 | 0.389 |
| reduced | 0.376 | -0.219 *** | 0.014 * | -0.030 ** | -- | 0.470 * | 9.872 (4.5) | 5.42e-6*** | 0.436 | 0.392 |
| 0 |  |  |  |  |  |  |  |  |  |  |
| full | -2.785 | -0.282 *** | 0.019 * | -0.044 ** | -0.009 | 0.610 * | 10.25 (5.5) | 8.42e-7 *** | 0.506 | 0.457 |
| reduced | -1.891 | -0.289 *** | 0.020 * | -0.048 ** | -- | 0.641 * | 12.63 (4.5) | 3.25e-7 *** | 0.498 | 0.458 |
| 6 |  |  |  |  |  |  |  |  |  |  |
| full | 2.738. | -0.117 * | -0.001 | -0.020 * | 0.005 | 0.119 | 3.192 (5.5) | 0.014 * | 0.242 | 0.166 |
| reduced | 3.358 *** | -0.093. | -- | -0.026 ** | -- | -- | 6.246 (2.6) | 0.004 ** | 0.182 | 0.153 |
| 12 |  |  |  |  |  |  |  |  |  |  |
| full | 2.618 ** | -0.088 ** | 0.005 . | -0.004 | -0.003 | 0.151 . | 3.419(5.5) | 0.010 ** | 0.255 | 0.18 |
| reduced | 2.267 ** | -0.087 ** | 0.005 | -- | -- | 0.155 . | 5.497 (3.5) | 0.002 ** | 0.241 | 0.197 |
| 24 |  |  |  |  |  |  |  |  |  |  |
| full | 3.384 *** | -0.0004 | 0.002 | -0.003 | 0.005 | 0.135 | 1.477 (5.5) | 0.214 | 0.129 | 0.041 |
| reduced | -- | -- | -- | -- | -- | -- | -- | -- | -- | -- |

Parameters, significance (F-statistics) and comparison of different models for total performance on IMRT and performance on separation conditions (0, 6, 12, 24 semitones). Models are compared using R<sup>2</sup> and R<sup>2</sup><sub>adjusted</sub>. Higher R<sup>2</sup> and more parsimonious models (fewer predictors) indicate superior models. Significance: p<.10 ., p<0.05 \*, p<0.01 \*\*, p<0.01 \*\*\* (uncorrected).

In summary, 39.2 % (R^2^_adjusted_) of variance on overall performance on IMRT and 45.8% (R^2^_adjusted_) of variance in the fully interleaved 0-ST-condition were explained by absolute pitch ability (ZMAD), musicality (MSI) and disembedding in vision (GEFT) (with IMRT version as a covariate). Even though R^2^_adjusted_ decreased for the easier conditions with a greater separation of probe and distractor melodies (6, 12), absolute pitch performance remained significant in both cases, while the other variables lost more of their influence (see table 4). Figure 4 shows regression models for overall performance (figure 4A) and separation conditions (figure 4B-E) with regression line corresponding to a simple linear regression of absolute pitch ability (Z_MAD) on d’ for simplicity reasons. Color and shape scales correspond to values on GEFT and MSI, respectively.

## Discussion

For the first time we investigated auditory streaming ability^1^ in absolute and relative pitch professional musicians using an interleaved melody recognition test^2,3^. Interestingly, general performance correlated with a visual embedded figures test (GEFT^47^, mean time needed to solve an item), which confirmed our hypothesis that the interleaved melody recognition test serves as an auditory hidden/embedded figures^6^. However, it must be made clear that direct measures of streaming ability using alternating high and low frequency tones^55^ may not necessarily have anything to do with the embedded figures concept. In the following we therefore stick to the terms “interleaved melody recognition test” and our acronym (IMRT) which we interpret to be a paradigm for both auditory streaming and the auditory embedded figures test.

General performance (sensitivity index d’, signal detection theory) as well as performance at different levels of frequency separation between probe and distractor melodies exceeded chance level (d’= 0) in both groups, pointing to a general advantage of professional musicians in auditory interleaved melody tests. This effect can be explained by above average musical knowledge in this sample which could either result in increased availability of musical schemas (schema-based auditory streaming^2^) or be based on heightened ability to recognize melodies per se. However, as we did not include a control condition in which no distractor tones were present in the second sequence of each trial, we cannot finally decide upon this issue. Ceiling effects obtained for the high separation condition (24-ST) are perfectly in line with previous reports of ceiling effects with respect to large distances between probe and distractor melodies^2^. Responses were only slightly biased towards “no” responses overall and this was most strongly the case for the most difficult 0-ST condition. As RP possessors always exhibited a more pronounced response bias this was seen as an indicator of uncertainty of responses, especially when the task was most challenging.

Furthermore, increasing the distance between probe and distractor melody (separation conditions) also increased performance of participants in both groups. Therefore and in accordance with other authors^2–4^ pitch separation as a perceptual cue seems to be of general importance for the processes of auditory streaming. However, in our study, we found an above chance level performance in the fully interleaved condition (0-ST). To our knowledge other authors have always found weak performance in healthy subjects for this condition and have argued for the necessity of this pre-attentive pitch cue based mechanism to come into play. In their view, schema-based auditory streaming might only be possible in interaction with bottom-up sensory processes (such as using pitch cues) that structure the auditory signal^2–4,6^. In line with this, increased performance in our sample of professional musicians may have been caused by extensive musical experience which led to additional musical schema-based processes in the 0-ST condition for these subjects. Musicians for example have daily exercise in extracting and focusing on melodies and fore vs. background in orchestral or other ensemble music as well as in solo performances of harmonical instruments like piano, which can play several melodies at the same time. This explanation would be consistent with evidence that auditory experience, such as familiarity with particular voices^56^ or words^57^, can influence the auditory streaming of speech. This hypothesis is further strengthened by the marginal correlation of Musical Sophistication Index (MSI)^43^ with overall performance and its contribution to some of the regression models. MSI is a general measure of musical sophistication aimed at the investigation of non-musicians which might explain the only small degree of correlation. It is possible that musicality tests which might be more suitable for trained musicians would have yielded higher correlations. However, as the interleaved melody recognition paradigm exhibits high similarity to famous tests of musicality (e.g. AMMA, Advanced measures of music audiation^46^)both in terms of stimuli and task (recognition of melodies), we did not include those tests.

The even higher performance of absolute pitch possessors in the fully interleaved and all other separation conditions as well as in overall performance allows us to assume that additional processes related to the absolute perception and/or naming ability of pitches lead to an advantage of AP’s in this test. Perhaps stable pitch-label associations comprise an additional schema-based mechanism, by which probe melody and tones in the interleaved sequence can be compared and extracted. However, as AP’s performance on IMRT also increases with the separation of probe and distractor melody, pitch-label associations might also increase auditory streaming ability due to an additional pre-attentive perceptual quality (pitch chroma) similar to e.g. tone-color perceptions of colored hearing synaesthets. Indeed, some authors have argued to explain absolute pitch ability as a form of synesthesia^31,58^, which also leads to enhanced low level perception^32,59^. Interestingly, enhanced low-level perceptional functioning respectively a tendency to focus on details are two main elements of famous theories to explain autistic symptoms^60–62^. Furthermore, nearly all special abilities up to savant skills in autism share certain features like enhanced low-level perception, focus on details and mapping of two cognitive or perceptual structures according to their inherent elemental structure (Theory of Veridical Mapping^31^). The one-to-one mapping of pitches to pitch labels might therefore be similar to the one-to-one mappings of pitches to colors or letters to colors. This could lead to enhanced performances on experiments in the affected sensory modalities. This enhancement might in turn be caused by pre-attentive processes that make use of the additional sensory quality that other subjects lack.

On top of that, autistic people can perform above change with fully interleaved sequences while faring relatively poorly with^29^. However, in our sample, none of the performances on IMRT was explained (even in part) by autistic traits. Despite that, we cannot rule out the possibility that the lack of correlation with autistic symptoms is due to very heterogeneous factors that play a role in the development of absolute pitch (e.g. heritability, onset of musical training, ethnicity). A tendency towards higher autistic traits in AP, as was also present in our sample, might form only one out of various different influences on the acquisition of AP. Therefore the lack of a correlation might be in part because the relation of AP and autistic traits is not true for all AP’s.

In general, performance on IMRT (sensitivity index d’), i.e. the ability to recognize a melody in an interleaved sequence was highly (R² = 15-39.2 %, smaller R² for easier conditions) dependent on absolute pitch proficiency as measured by a pitch adjustment test (PAT, developed after Dohn et al.^48^), visual disembedding (GEFT) ^47^ and in the more difficult conditions musical sophistication (MSI)^43^. We therefore conclude that the interleaved melody recognition test is an auditory streaming paradigm^5,6^ which additionally measures the general ability to find auditory hidden figures (as for the correlation with GEFT) and that performance on it is enhanced by musical training and absolute pitch ability. The latter might be due to an additional sensory quality leading to enhanced low-level perceptual functioning in general. This study therefore to our knowledge is the first to show enhanced auditory disembedding in absolute pitch possessors. Nevertheless some constraints have to be mentioned: First, we did not include a control condition for melody recognition (see above), second, due to a technical error the number of trials per condition and subject was not counterbalanced (see Methods section). However, additional statistical analyses indicated that results were not affected by this issue (see Methods section and Supplementary Material A). Third, with respect to the discussed similarities to autism, a third subsample including autistic people would have been desirable.

To conclude we would like to draw attention to goals that future studies should address. Because of the partial, but nevertheless re-occurring, similarities between absolute pitch possessors and autism in terms of cognition, perception, personality traits and neurophysiology and anatomy it remains to be investigated whether and to what extend one is the cause or a side effect of the other, or which external factors lead to the coincidence. Such investigation would not only increase the knowledge about both phenomena but also help to understand fine-graded differences in human perception and its relation to other cognitive functions and personality traits.

## Acknowledgements

We thank Dr. C. Ioannou for assistence in creating the figures. We further want to thank Hannes Schmidt, Pablo Carra, Artur Ehle, Fynn Lautenschläger, Michael Großbach, Christos Ioannou, Daniel Scholz and all other colleagues for fruitful discussions and technical assistance.

## Authors Contributions Statement

TW designed the study, collected, analysed and interpreted the data. EA contributed to the design of the study and interpretation of the data. Y.H. took part in data collection and analysis. All authors read, improved and approved the final manuscript.

## Additional Information

### Ethics approval and consent to participate

The study was approved by the ethic committee of the Hanover Medical School (Approval no. 7372, committee’s reference number: DE 9515). All participants gave written consent.

### Availability of data and material

The datasets generated and/ or analysed during the current study are not publicly available due to specifications on data availablity within ethics approval. Data are however available from the corresponding author upon reasonable request and with permission of the ethics committee of the Hanover Medical School.

